# A novel enhancer blocker assay identifies AT-rich simple sequence repeats as D1-dependent enhancer blockers in *Drosophila*

**DOI:** 10.1101/2024.12.04.626877

**Authors:** Kundurthi Phanindhar, Fathima Athar, Rakesh K Mishra

**Author notes:** Corresponding authors: Rakesh K Mishra, Kundurthi Phanindhar.

## Abstract

Simple sequence repeats (SSRs) are tandem repetitions of 1-6 bp DNA motifs at least 12 bp long. Certain length-enriched SSRs function as enhancers/silencers, enhancer blockers and barriers in a mammalian cell line. However, whether SSRs have similar cis-regulatory functions *in vivo* and their underlying mechanisms are unknown. To address this, we looked for SSR-binding proteins and investigated if SSRs function as enhancer blockers *in vivo*. We developed a novel *Drosophila* assay to assess enhancer blocker activity *in vivo* and to circumvent the low throughput and position effects seen in traditional enhancer blocker assays. Our assay uses endogenous *vestigial* gene enhancers, which, when blocked, result in easily scorable wing phenotypes increasing throughput. The attP-attB-based recombination system to integrate test fragments at specific site avoids positional effects. Furthermore, using EMSAs and DNA pull-downs followed by LC-MS/MS, we show that SSRs bind to proteins in a sequence-specific manner and identify 33 unique SSR-binding proteins. One of these proteins, D1, was enriched in several SSRs, viz., AAT_14_, AAAT_10_, AAAAT_8_, AATAT_9_, AAAG_13_ and AAAAG_11_. Using our novel *vestigial in vivo* enhancer blocker assay, we show for the first time that AT-rich SSRs function as enhancer blockers in a D1-dependent manner in *Drosophila*.

## Introduction

Simple sequence repeats (SSRs), microsatellites or simple tandem repeats (STRs) are tandem repetitions of 1-6 bp DNA motifs with a minimum size of 12 bp (Subramanian et al. 2003). The 1-6 bp DNA motifs can have 5356 sequence variations, which are grouped into 501 SSR classes (Avvaru et al. 2017). SSRs function across bacteria to humans, regulate many biological processes including transcription (Hammock and Young 2005; Horton et al. 2023; Staib et al. 2002; Vinces et al. 2009; Xu et al. 1997) and splicing (Hefferon et al. 2004; Hui et al. 2003, 2005), and are used as genetic markers (Dietrich et al. 1992; Jacob et al. 1991; Litt and Luty 1989; Weber and May 1989). Though long stretches of SSRs are less likely, certain SSRs are enriched in the genomes at longer lengths, indicating specific length-dependent functions (Ramamoorthy et al. 2014; Srivastava et al. 2019). A cis-regulatory function was demonstrated for the 23 length-enriched SSRs in the human genome in a mammalian cell line where SSRs functioned as enhancers or silencers, enhancer blockers and barriers (Krishnan et al. 2017). However, the basis of the cis-regulatory functions is unknown, and the cis-regulatory functions were not confirmed *in vivo*, in an animal model.

SSRs like AGAT and GA function as enhancer blockers (Kumar et al. 2013; Vasanthi et al. 2010), a boundary element that prevents the interaction between an enhancer and its promoter only when present between them (Geyer and Corces 1992). These repeats have been shown to have protein-binding sites. For example, CLAMP and GAF bind to GA-rich repeats (Kaye et al. 2018). GAF is known to regulate the enhancer blocker activity of GA repeats (Vasanthi et al. 2010). GAF has one C2H2 zinc finger and binds both long and short GA repeats. On the other hand, CLAMP has 6 C2H2 zinc fingers that bind only long GA repeats (Kaye et al., 2018). Both CLAMP and GAF are shown to regulate the Fab-7 boundary activity (Kaye et al. 2017). Since several length-enriched SSRs were shown previously to have enhancer blocker function in the cell culture system (Krishnan et al. 2017), we sought to identify the proteins binding to these SSRs and test if the length-enriched SSRs function as enhancer blockers *in vivo*.

Traditionally, enhancer blocker assays in *Drosophila* have been P-element based. The P-elements are integrated randomly into the genome and are subject to position effects, making it difficult to compare test elements. Moreover, the P-element based assays depend on reporter genes such as *mini-white* or *lacZ*, which require molecular biological assays for quantification, and hence are low throughput (Kellum and Schedl 1992; Vazquez and Schedl 1994; Hagstrom et al. 1996; Cai and Levine 1995).

In the present study, we show that several length-enriched SSRs bind proteins in a sequence-specific manner. We identify proteins that are unique to a few SSRs, and those that bind multiple SSRs. To circumvent the problems of traditionally used enhancer blocker assays, we developed a novel *in vivo vestigial* enhancer blocker assay in *Drosophila,* which utilizes the endogenous *vestigial* gene locus where the test (SSRs) DNA fragments are integrated in a site-directed fashion. Blocking the *vestigial* gene enhancers by an enhancer blocker produces an easily scorable morphological phenotype, making the assay high throughput. Using our *in vivo vestigial* enhancer blocker assay, we show that AT-rich SSRs and their binding protein D1 act as enhancer blockers in *Drosophila*.

## Results

### 1. *Drosophila* embryo nuclear extracts have SSR-binding proteins

SSRs like AGAT and GA function as enhancer blockers in human cell lines and *Drosophila* (Kumar et al. 2013; Vasanthi et al. 2010). Moreover, the pathology and the proteins involved in human repeat expansion disorders are conserved in *Drosophila* (Miller et al. 2000; Warrick et al. 1998; Zhang et al. 2015). Thus, *Drosophila* is a good model system to investigate the functions of SSRs. We used 0-16 h *Drosophila* embryo nuclear extracts to identify SSR-binding proteins. We used SSRs that were earlier shown to be enriched in the human genome at longer lengths than expected and to have cis-regulatory functions (Ramamoorthy et al. 2014; Krishnan et al. 2017). Double-stranded DNA oligos corresponding to the SSRs were end-labelled and protein binding was tested using Electrophoretic mobility shift assays (EMSA).

All 15 SSRs tested showed gel retardation in EMSA suggesting that SSRs bound nuclear proteins from 0-16 h *Drosophila* embryos (Figure 1A-H and Sup. Figure 1A-G). We proceeded to confirm the specificity of these DNA-protein complexes (EMSA shifts) by competition with SSRs identical, similar, or dissimilar to the test SSR. SSRs were considered similar if the sequences were 75-80% identical and dissimilar if they were 25-50% identical. We performed EMSAs with 15 different SSRs and describe one in detail here. Radiolabeled test SSR AAT_14_ bound proteins in the nuclear extract, indicated by the mobility shift in EMSA (Figure 1A, lane 2). AAT_14_-DNA-protein complexes were competed out by the identical unlabeled AAT_14_ added at 10, 20, or 50-fold higher concentration than radiolabeled AAT_14_. This is evident in the disappearance of the shift and release of free radiolabeled AAT_14_ probe (Figure 1A, lanes 3-5). However, SSRs dissimilar to AAT_14_, such as AAGG_11_ or ATCC_9_, even at a 50-fold higher concentration, do not affect the shift of radiolabeled AAT_14_ DNA-protein complexes (Figure 1A, lanes 8 and 11). This indicated that proteins in 0-16 h *Drosophila* embryo nuclear extracts bind AAT_14_ in a sequence-specific manner.

**Figure 1:**
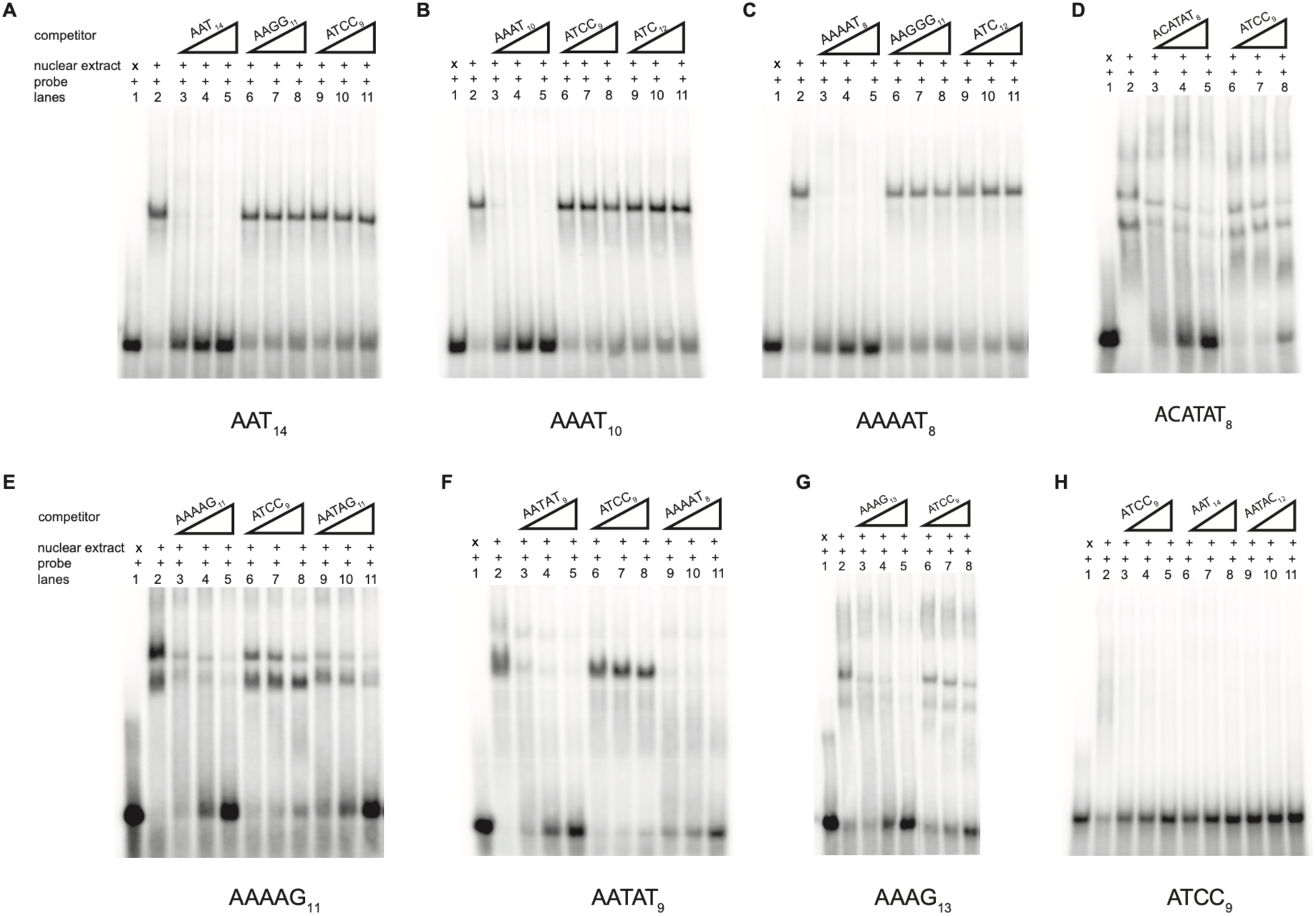
Electrophoretic mobility shift assays of length-enriched SSRs in the presence of nuclear extract from 0-16 h Drosophila embryos. **A-H)** Each gel shows DNA-protein complexes of radiolabeled SSR in lane 2. Competition reactions with the same but unlabeled SSR are shown at 10- or 20- or 50-fold concentrations higher than radiolabeled probe in lanes 3-5. Competition reactions with a different unlabeled SSR (dissimilar in sequence) to the radiolabeled probe are shown at 10- or 20- or 50-fold concentrations in A-H- lanes 6-8; A-C, H -lanes 9-11. Competition reactions with a different unlabeled SSR with sequence similarity to the radiolabeled probes are shown at 10- or 20- or 50-fold concentrations in lanes 9-11 of gels E, F.

Some SSRs tested, such as ACATAT_8_, AAAAG_11_, AAAG_13_ and AATAG_11_ showed multiple gel shifts in EMSAs, indicating the presence of either multiple SSR-binding proteins for a single SSR or that the binding protein forms multimeric complexes (Figure 1D-E, G and Sup. Figure 1D). Alignment of stretches of SSRs showed that some SSRs share motifs that are either identical or differ by one or a few nucleotides. For example, the repeating motif in AAAAT_8_, ‘AAAAT’, differs from the motif of AATAT_9_, ‘AATAT’, by one nucleotide in the third position. Accordingly, proteins binding to AATAT_9_ were competed out by AAAAT_8_ (Figure 1F, lanes 9-11). Likewise, AATAG_11_-DNA-protein complexes were competed out by AATAC_12_ (Sup. Figure 1D, lanes 9-11) and similar results were also observed with AAAAG_11_ (Figure 1E, lanes 9-11) and AAGAG_12_ (Sup. Figure 1C, lanes 9-11). This indicates that more than one SSR may bind the same protein(s). ATCC_9_ showed a gel shift in our binding conditions, which was easily competed out by dissimilar SSRs such as AAT_14_ and AATAC_12_ used as competitors, suggesting a lack of specificity between ATCC_9_ and the proteins bound (Figure 1H). Overall, we found that 14 of the 15 SSRs we tested bound proteins in the *Drosophila* embryo (0-16 h) nuclear extract in a sequence-specific manner (Figure 1A-G and Sup. Figure 1A-G).

### 2. Identification of *Drosophila* SSR-binding proteins using proteomics

We chose eight SSRs, seven of which - AAT_14_, AAAT_10_, AAAAT_8_, AAAG_13_, AAAAG_11_, ACATAT_8_ and AATAT_9_ showed distinct shifts and sequence-specific protein binding in EMSAs with nuclear extract from 0-16 h *Drosophila* embryos in our experiments (Figure 1A-G). The eighth SSR AGAT_10_ was previously shown to bind proteins in an EMSA assay and function as an enhancer blocker (Kumar et al. 2013). Among the seven, AAT_14_, AAAT_10_ and AAAAT_8_ showed a single shift (Figure 1A-C), AAAAG_11_ and AAAG_13_ showed two shifts (Figure 1E and 1G), and ACATAT_8_ showed multiple shifts in their EMSAs (Figure 1D). AATAT_9_ was previously shown to bind to D1 protein and hence was included as a positive control for the biotinylated DNA-based protein pulldown assay (Levinger 1985).

To isolate SSR-binding proteins, biotinylated DNA oligos were used in binding reactions with nuclear extracts from *Drosophila* embryos (0-16 h) and the isolated SSR-binding proteins were identified using LC-MS/MS (Figure 2A). We identified 1320 proteins after eliminating potential contaminants, peptides identified only by site, reverse hits and proteins identified by only one unique peptide. The percentage of proteins identified in at least two biological replicates across SSRs was between 80.67% to 91% (Figure 2B).

**Figure 2:**
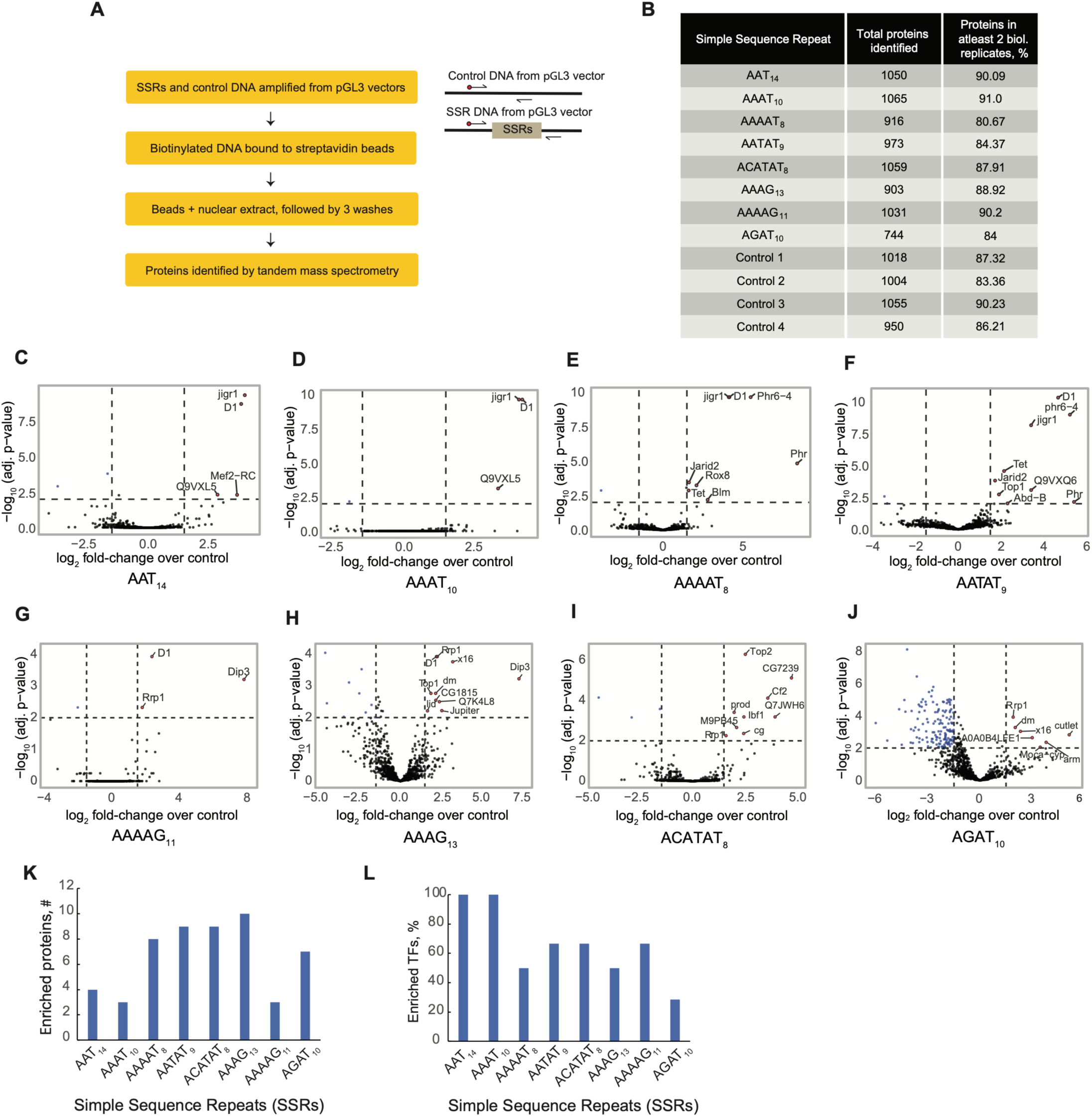
Proteomics of SSR associated proteins. **A)** Flowchart depicting the strategy used for isolation of SSR binding proteins and schematic representing the amplification of biotinylated DNA from *pGL3* vectors. **B)** Table showing the number and percentage of proteins pulled down across various SSR pull-downs. **C-J)** Volcano plots showing differentially enriched proteins (red dots with gene names) across SSR pull-downs of AAT_14,_ AAAT_10,_ AAAAT_8,_ AATAT_9,_ AAAAG_11,_ AAAG_13,_ ACATAT_8_ and AGAT_10._ The criteria for enrichment: logz(LFQ intensity fold change over control)> 1.5 and -log_10_(adj. p-value) > 2. **K)** Plot showing the number of enriched proteins obtained in the biotin labelled SSR pull-downs with *Drosophila* embryo (0-16 h) nuclear extract. **L)** Plot showing the percentage of enriched Transcription factors obtained in the biotin labelled SSR pull-downs with *Drosophila* embryo (0-16 h) nuclear extract.

Differentially enriched proteins between SSRs and their respective controls were identified using LFQ analyst with default parameters (Shah et al. 2020). We narrowed down proteins whose log_2_ (LFQ intensity fold change over control) was > 1.5 and -log_10_ (adj. p-value) was > 2 (Figure 2C-J). The number of proteins identified for each SSR were as follows: AAAAG_11_ - 3, AAAAT_8_ - 8, AAAT_10_ - 3, AAT_14_ - 4, AATAT_9_ - 9, ACATAT_8_ - 9, AGAT_10_ - 7, AAAG_13_ - 10 (Sup. Table 1 and Figure 2K). Several of these proteins bound multiple SSRs. AATAT SSR-binding D1 protein was identified among the proteins enriched with AATAT_9_. Several proteins enriched in the ACATAT_8_ pulldown, like Combgap, CG7239, CG1888 (Q7JWH6), prod and CG14005 (M9PB45), have been reported to be immunoprecipitated by Combgap (Chetverina et al. 2022). These observations confirmed the validity of our DNA-pulldowns and the analysis parameters.

We observed that several proteins bound more than one SSR, whereas D1, Jigr1 and Rrp1 bound multiple SSRs (Sup. Table 2). Further, six proteins commonly enriched between AAAAT_8_ and AATAT_9_ (Figure 2E-F) whose repeat motifs are 80% similar and AAAAT_8_ had competed out the protein binding to AATAT_9_ in EMSA (Figure 1F). D1 was enriched in 6 SSRs-AAT_14_, AAAT_10_, AAAAT_8_, AATAT_9_, AAAG_13_ and AAAAG_11_, where in SSRs AAAAT_8_ and AAAAG_11_, and AAAT_10_ and AAAG_13_ motifs differed by one nucleotide (Figure 2C-F, G-H). D1 was ∼2.5-fold higher enriched in AAAAT_8_ compared to AAAAG_11_, whereas it was ∼3.5-fold enriched in AAAT_10_ compared to AAAG_13_. However, Dip3 was the most enriched protein in AAAG_13_ and AAAAG_11_, with enrichment over AAAT_10_ and AAAAT_8_ was 256-fold and 128-fold, respectively (Figure 2G-H and Sup. file 1). Jigr1 was enriched in AAT_14_, AAAT_10_, AAAAT_8_ and AATAT_9_ (Figure 2C-F). Rrp1 was enriched in AAAG_13_, AAAAG_11_, ACATAT_8_ and AGAT_10_ (Figure 2G-J).

Since AAT_14_, AAAT_10_, AAAAT_8_ and AATAT_9_ have common proteins such as D1 and Jigr1 binding to them. We aligned these SSRs and observed a common motif “AATAA” recurring among the SSRs AAT_14_, AAAT_10_, AAAAT_8_ (Sup. Figure 2A). The recurring motif “AATAA” differs from “AATAT”, the motif in AATAT_9_ by a single nucleotide (Sup. Figure 2A). Since, AAT_14_, AAAT_10_, AAAAT_8_ and AATAT_9_ have a recurring motif, we hypothesized that protein binding to any of these SSRs should be competed out by the other SSRs. Thus, we used competition EMSAs to test the protein binding to radiolabeled AAAT_10_, which showed a shift with *Drosophila* 0-16 h embryo nuclear extract. This shift in AAAT_10_ was competed out with higher concentrations of the SSRs AAT_14_, AAAAT_8_, AAAT_10_ and AATAT_9_ (Sup. Figure 2B and 2E). We also show that protein binding to radiolabelled AAT_14_ or AAAAT_8_ is competed out by AAT_14_, AAAT_10_ and AAAAT_8_ indicating that common proteins bind the SSRs AAT_14_, AAAT_10_ and AAAAT_8_ (Sup. Figure 2F-G). We also observed that the protein binding to AAAT_10_ is competed to a lesser degree by AAAAG_11_ and AAAG_13_ (Sup. Figure 2C). However, other SSRs, including ACATAT_8_, AGAT_10_ and ATC_12_ did not compete out the protein binding to AAAT_10_ (Sup. Figure 2D-E). Overall, this suggests that common proteins may bind the AT-rich SSRs, AAT_14_, AAAT_10_, AAAAT_8_ and AATAT_9_, which may include D1 and Jigr. The LFQ intensities of D1 rather than Jigr1 across various SSR pull-downs match the competition pattern seen in AAAT_10_ competition EMSAs (Sup. Figure 2H-I).

### 3. Transcription factors, epigenetic regulators and DNA repair proteins are enriched in SSR-binding proteins

SSRs are known to regulate gene transcription (Albanèse et al. 2001; Horton et al. 2023; Gebhardt et al. 1999), are enriched in heterochromatin (Cuadrado and Jouve 2011; Lohe et al. 1993; Pimpinelli et al. 1995), have a role in RNA splicing (Hui et al. 2005, 2003; Hefferon et al. 2004) and may be involved in epigenetic regulation (Gymrek et al. 2016).

Of the 53 SSR-binding proteins, we identified 19 unique known or predicted transcription factors (TFs) across 8 SSR pull-downs. All 8 SSRs bound at least one TF (Table 1). Percentage enrichment of TFs for each SSR is shown in Figure 2L. Of the 19 unique TFs, 12 TFs bound exclusively to one SSR, while 7 bound to >1 SSR. The DNA binding domains in these 19 SSR binding TFs included MADF (myb/SANT-like domain in Adf-1), zinc finger domain, ARID (AT-rich interaction domain) domains, AT-hook, BED finger, basic helix-loop-helix (bHLH), homeobox, MADS (MCM1, AG, DEFA and SRF)-box and pipsqueak-type helix-turn-helix (HTH) domain (Table 1). The most common MADF domain was observed in 5 TFs, 4 TFs had the zinc finger domains and 1 TF had both MADF domain and zinc fingers (Table 1).

**Table 1:**
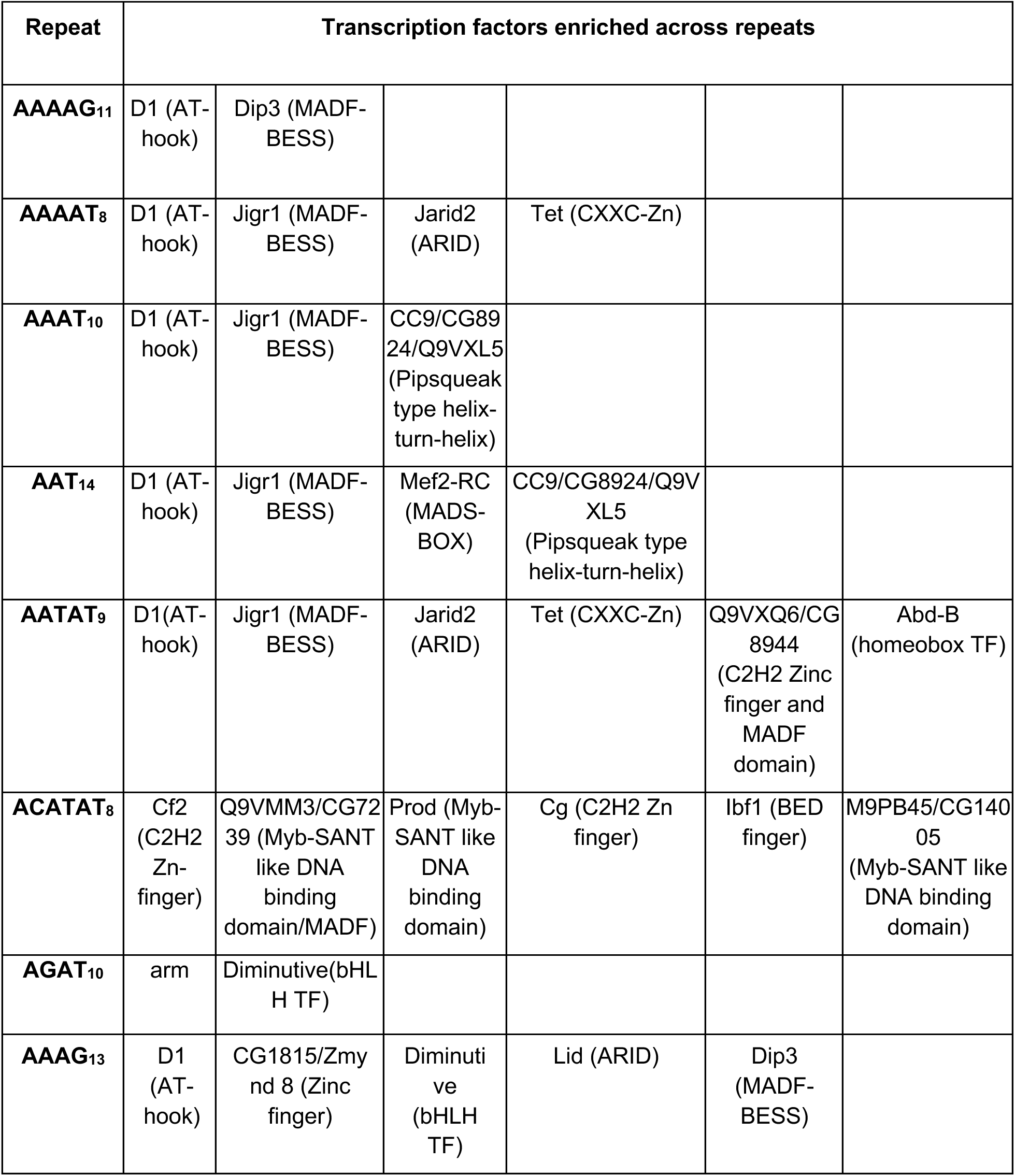
List of enriched Transcription factors enriched across various repeats along with their DNA binding domain indicated in brackets.

| Repeat | Transcription factors enriched across repeats |  |  |  |  |  |
| --- | --- | --- | --- | --- | --- | --- |
| **AAAAG<sub>11</sub>** | D1 (AT-hook) | Dip3 (MADF-BESS) |  |  |  |  |
| **AAAAT<sub>8</sub>** | D1 (AT-hook) | Jigr1 (MADF-BESS) | Jarid2 (ARID) | Tet (CXXC-Zn) |  |  |
| **AAAT<sub>10</sub>** | D1 (AT-hook) | Jigr1 (MADF-BESS) | CC9/CG8924/Q9VXL5 (Pipsqueak type helix-turn-helix) |  |  |  |
| **AAT<sub>14</sub>** | D1 (AT-hook) | Jigr1 (MADF-BESS) | Mef2-RC (MADS-BOX) | CC9/CG8924/Q9VXL5 (Pipsqueak type helix-turn-helix) |  |  |
| **AATAT<sub>9</sub>** | D1(AT-hook) | Jigr1 (MADF-BESS) | Jarid2 (ARID) | Tet (CXXC-Zn) | Q9VXQ6/CG8944 (C2H2 Zinc finger and MADF domain) | Abd-B (homeobox TF) |
| **ACATAT<sub>8</sub>** | Cf2 (C2H2 Zn-finger) | Q9VMM3/CG7239 (Myb-SANT like DNA binding domain/MADF) | Prod (Myb-SANT like DNA binding domain) | Cg (C2H2 Zn finger) | lbf1 (BED finger) | M9PB45/CG14005 (Myb-SANT like DNA binding domain) |
| **AGAT<sub>10</sub>** | arm | Diminutive(bHLH TF) |  |  |  |  |
| **AAAG<sub>13</sub>** | D1 (AT-hook) | CG1815/Zmynd 8 (Zinc finger) | Diminutive (bHLH TF) | Lid (ARID) | Dip3 (MADF-BESS) |  |

Many proteins associated with the heterochromatin, such as D1, Prod, Topoisomerase 2, CG1815 and Rrp1 were identified in our SSR pull-downs. D1 protein bound to AT-rich SSRs such as AAT_14_, AAAT_10_, AAAAT_8_, AAAG_13_ and AAAAG_11_ in addition to AATAT_9_. D1, an AT-hook protein, is responsible for chromocenter maintenance in spermatogonial cells and has a role in heterochromatin maintenance, exemplified by its effect on the *w[m4h]* PEV model (Jagannathan et al. 2018; Aulner et al. 2002). Prod (proliferation disrupter), a MADF domain-containing protein and Topoisomerase 2 bound ACATAT_8_. Prod is shown to be responsible for chromocenter formation in imaginal discs and lymph glands, like the D1 protein (Jagannathan et al. 2019), and Topoisomerase 2 is known to contribute to heterochromatin maintenance (Blattes et al. 2006; Napoletano et al. 2021). CG1815 (Zmnyd 8), a zinc finger-containing protein and an interactor of HP1a (Alekseyenko et al. 2014), bound to AAAG_13_ SSR. Rrp1 (Recombination repair protein 1), a protein associated with both right and left telomere-associated repeats (Antão et al. 2012) bound to ACATAT_8_, AAAG_13_, AAAAG_11_ and AGAT_10_ SSRs.

Several proteins involved in polycomb regulation bound to 5 SSRs used in our assay. Jigr1 (Jing interacting gene regulatory 1), a MADF-containing transcription factor, a known regulator of domino (Ellis et al. 2015), and interactor of Polycomb (Pc) (Kang et al. 2015), bound to AAT_14_, AAAT_10_, AAAAT_8_ and AATAT_9_. E(z) (Enhancer of zeste)-interacting protein, Jarid2 (Jumonji, AT-rich interactive domain 2) (Kang et al. 2015) bound to AAAAT_8_ and AATAT_9_ SSRs. Combgap (Cg), another protein known to recruit polycomb complexes to PREs, was enriched in proteins binding to ACATAT_8_ SSR (Ray et al. 2016). Chorion factor 2(Cf2), which is a paralog of Cg, bound to ACATAT_8_ and has been shown to bind to the sequence GTATATTAT, where GTATAT is the reverse complement of ACATAT (Hsu et al. 1992).

RNA binding proteins, Rox8 and X16 (x16 splicing factor), regulate splicing (Bradley et al. 2015; Park et al. 2004). Rox8 was enriched among the proteins bound to AAAAT_8_, while X16 was enriched in proteins binding to AAAG_13_ and AGAT_10_. Among the DNA repair proteins, Phr (Photorepair) and Phr6-4 ((6-4)-Photolyase), which help repair DNA damage caused by UV exposure (Boyd and Harris 1987; Todo et al. 1996), bound to AAAAT_8_ and AATAT_9_. Rrp1, involved in base excision and recombination-mediated DNA repair (Takeuchi et al. 2006; Sander et al. 1991), was enriched in proteins binding to AAAG_13_, AAAAG_11_, ACATAT_8_ and AGAT_10_ SSRs.

Further, many uncharacterized SSR-binding proteins CG8924 (Q9VXL5), CG8944 (Q9VXQ6), CG1888, CG7239, CG7878 (Q7K4L8), CG14005 and CG10139 (A0A0B4LFE1) were also identified in our DNA-based protein pulldown assay. While molecular details of how SSRs are involved in RNA splicing, transcription and DNA repair are not well known, the protein classes identified are in line with functions previously ascribed to SSRs (Albanèse et al. 2001; Horton et al. 2023; Cuadrado and Jouve 2011; Lohe et al. 1993; Hui et al. 2003; Hefferon et al. 2004; Gymrek et al. 2016).

### 4. Development and characterization of a novel enhancer blocker assay in *Drosophila*

D1 protein was highly enriched in the proteomic pull downs with 4 AT-rich SSRs namely AAT_14_, AAAT_10_, AAAAT_8_ and AATAT_9_; We next investigated if D1, along with these AT-rich SSRs, functions as enhancer blockers *in vivo*. Traditionally, enhancer blocker assays in *Drosophila* are P-element transposon-based and subject to position effects as they integrate in the genome at random loci. The assays have low throughput, requiring molecular biology-based quantifications (Kellum and Schedl 1992; Vazquez and Schedl 1994; Hagstrom et al. 1996; Cai and Levine 1995). To address these drawbacks, we developed a novel enhancer blocker assay which allows site-specific integration of test fragments using the attP-attB based recombination system, thereby avoiding non-specific position effects (Bateman and Wu 2008; Groth et al. 2004). The assay utilizes the endogenous enhancers of the *vestigial* gene, which, when blocked, result in wing abnormalities, allowing direct and visible quantification.

*vestigial* gene (*vg*) is essential for *Drosophila* wing and haltere development and is regulated by the D/V-enhancer and the quadrant enhancer. D/V-enhancer drives *vg* expression in the dorsoventral (DV) and anteroposterior (AP) margins of the wing pouch. In contrast, the quadrant enhancer drives the *vg* expression in the wing pouch quadrants demarcated by DV and AP margins (Figure 3A) (Williams et al. 1991; Kim et al. 1996). D/V-enhancer deletion results in complete loss of wings and halteres, as exemplified in *vg[83b27]* (Williams et al. 1991). Whereas, deletion of the quadrant enhancer was recently shown to be dispensable for wing development (Farfán-Pira et al. 2022). We expected that blocking the D/V-enhancer with an enhancer blocker should result in the loss of wings and halteres.

**Figure 3:**
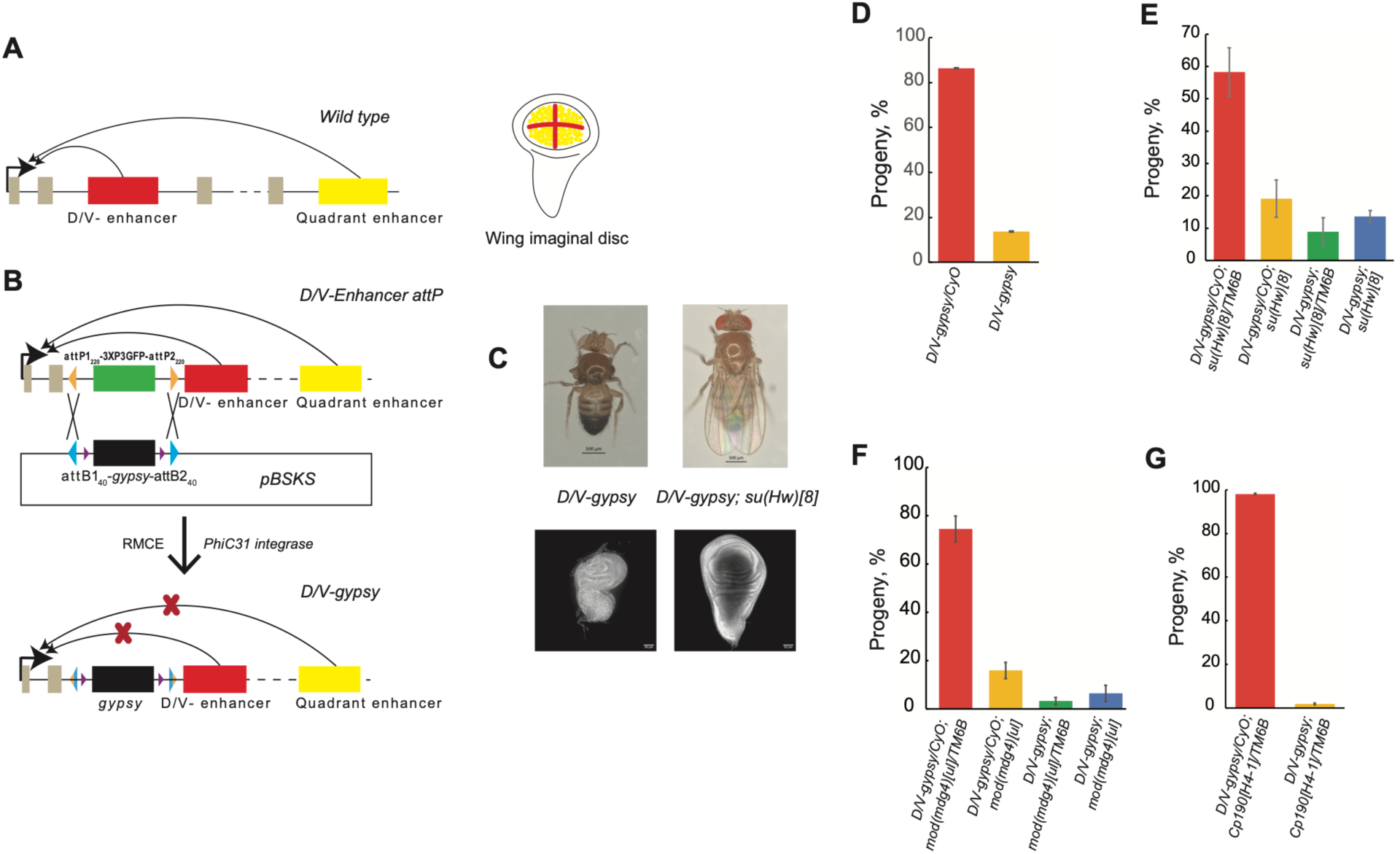
Development of novel *vestigial* enhancer blocker assay. **A)** Schematic depicting the *vestigial* locus with its well-characterized ON-enhancer (red box) and quadrant enhancer in wild type (yellow box). The wing imaginal disc on the right shows the areas of the wing pouch where the ON-enhancer (red lines) and quadrant enhancer (yellow dots) are active. **B)** Schematic of recombination between the *attP_220_*sites of *ON-enhancer attP* line and the *attB_40_* sites of *pBSKS* donor plasmid containing *attB1_40_-gypsy-attB2_40._* The *ON-enhancer attP* line has *attP1_220_-3XP3GFP-attP2_220_* insertion upstream of the ON-enhancer (red box). The *attP_220_*sites (orange triangles) and *3XP3GFP* (green rectangle) are represented. *gypsy* (black rectangle) and *att8_40_* sites (blue triangles) are depicted in the *pBSKS* donor plasmid. The plasmid also harbors *loxP* sites (violet triangles) for *gypsy* excision. The recombination between *attP_220_* and *attB_40_* sites results in RMCE, which ultimately leads to GFP loss in the *ON-gypsy* line. **C)** Representative images of homozygous *ON-gypsy* fly showing absence of wings and halteres (top left) and OAPI stained wing imaginal discs from the 3rd instar larvae of the same genotype showing abnormal morphology (bottom left). *ON-gypsy* homozygous fly homozygous for *su(Hw)* mutation shows the rescue of wings (top right) and OAPI stained wing imaginal discs from the 3rd instar larvae of the same genotype showing normal morphology (bottom right). **D)** Percentage of progenies obtained on selfing *ON-gypsy/CyO* flies. **E)** Percentage of progenies obtained on selfing *ON-gypsy/CyO; su(Hw)[B]ITM6B* flies. **F)** Percentage of progenies obtained on selfing *ON-gypsy/CyO; mod(mdg4)[ul]ITM6B* flies. **G)** Percentage of progenies obtained on selfing *ON-gypsy/CyO; Cp190[H4-1]/TM6B* flies.

We generated two *attP* lines with the *attP* landing sites (*attP1_220_-3XP3GFP-attP2_220_*) inserted in the 2nd intron of the *vestigial* gene using the CRISPR-Cas9 system. In the *D/V-enhancer attP* line, the *attP* landing site was upstream of the D/V-enhancer, whereas the *quadrant-enhancer attP* line had the *attP* landing site downstream of the D/V-enhancer. The *attP* landing site insertion in the *vg* locus results in GFP expression in the eyes (Sup. Figure 3A). The *D/V-enhancer attP* line assays for blockade of both the D/V-enhancer and quadrant enhancer, whereas *quadrant attP* line assays the blockade of only the quadrant enhancer. Both the *attP* lines have normal wings and halteres, suggesting that the *attP* landing site insertions did not affect any functional elements. To accomplish recombination-mediated cassette exchange (RMCE) between the *attP_220_* sites of the *vestigial* locus and the *attB_40_* sites of the *attB* donor plasmid (*pBSKS_attB1_40_-*

*test element-attB2_40_*), the *attP* transgenic lines were brought in the background of *nanos-phiC31 integrase*. Upon RMCE, the *3XP3GFP* in the *vestigial* locus is exchanged with test DNA from the donor plasmid (Figure 3B). Thus, flies in which RMCE was successful show loss of GFP in the eyes (Sup. Figure 3B).

### 5. *gypsy* functions as an enhancer blocker in our *vestigial* enhancer blocker assay

We used *gypsy*, one of the strongest known enhancer blockers, as a positive control to test the *vestigial* enhancer blocker assay. *gypsy* integrated *D/V-enhancer attP* line referred to as *D/V-gypsy* showed ∼60% lethality in homozygous conditions. When heterozygous *D/V-gypsy* lines were selfed, only 13.7% of homozygous flies survived as opposed to the expected 33% (Figure 3D). Among the surviving homozygous flies, 16.67% had no wings, ∼41.67% had one wing and the rest 41.67% of flies had normal wings (Figure 3C and Sup. Figure 3C). Almost all homozygous flies show upright post-scutellar bristles and around 80% show the absence of both halteres (Figure 3C and Sup. Figure 3D). 80% of the flies that had one wing also show a transformation to notum (Sup. Figure 3D red box). All these phenotypes are characteristic of *vestigial* mutants (Lindsley et al. 1967; Williams and Bell 1988; Bownes and Roberts 1981; Williams et al. 1990).

*gypsy* integration in the *quadrant-enhancer attP* line, which blocks only the quadrant enhancer, did not result in any phenotype. Quadrant enhancer has been shown to be dispensable for wing development (Farfán-Pira et al. 2022). This also suggests that the loss of wings in *D/V-gypsy* flies may not be due to epigenetic repression of the *vestigial* enhancers. We used the *D/V-enhancer attP* line for further experiments.

A protein complex consisting of proteins Su(Hw), Mod(mdg4) and Cp190 is known to bind and regulate *gypsy* enhancer blocker activity (Gerasimova et al. 1995; Pai et al. 2004; Spana et al. 1988). We brought *D/V-gypsy* lines in the mutant background of either *su(Hw)*, *mod(mdg4)*, or *Cp190*. *D/V-gypsy* heterozygous flies had normal wings and halteres irrespective of the *su(Hw)* mutant dosage. Meanwhile, the *D/V-gypsy* homozygous flies showed a rescue in lethality, with 13.5% of flies surviving against the expected 11.11% upon homozygous *su(Hw)* mutation. These flies also had normal wings (5% flies showed mild notches) and halteres (Figure 3C and 3E). As expected, the homozygous *su(Hw)* mutation rescued the abnormal morphology of the wing imaginal disc from the 3rd instar larvae of homozygous *D/V-gypsy* flies (Figure 3C). However, 80% of homozygous *D/V-gypsy* flies had no halteres and 13% had defective wings upon heterozygous *su(Hw)* mutation. Thus, *su(Hw)* mutation dosage dictates the amount of phenotype rescue. In contrast, only 6% *D/V-gypsy* flies were homozygous when brought in the homozygous mutant background of *mod(mdg4),* indicating partial rescue (Figure 3F). Additionally, only 2% *D/V-gypsy* homozygous flies were observed when brought in the mutant background of *Cp190,* probably because the homozygous *Cp190* mutation is lethal (Figure 3G). The partial rescue by *mod(mdg4)* and *Cp190* could also be because *su(Hw)* is the major regulator of *gypsy* function. Collectively, these results validate our novel *vestigial* enhancer blocker assay by demonstrating that *gypsy* functions as an enhancer blocker in *Drosophila*.

### 6. AT-rich SSRs function as enhancer blockers in *Drosophila*

AGAT and GA SSRs were shown to function as enhancer blockers in *Drosophila* and human cells (Kumar et al. 2013; Vasanthi et al. 2010). A previous study in K562 cells showed that length-enriched SSRs have cis-regulatory functions (Krishnan et al. 2017). So, we sought to investigate if some of these SSRs show similar function *in vivo*, and understand their mechanism of action. Since we identified that D1 binds better to AAT_14_, AAAT_10_, AAAAT_8_ and AATAT_9_ we integrated these SSRs into the *D/V enhancer attP* line and generated four different ‘*D/V-SSR*’ lines. We assayed wing phenotypes in multiple homozygous *D/V-SSR* lines for each SSR and observed normal wings except for a small percentage of flies that showed a mild phenotype (Sup. Figure 3E) similar to T2 phenotype described below-*D/V-AAT_14_* - 0.88%, *D/V-AAAT_10_* - 0.43%, *D/V-AAAAT_8_* - 1.15% and *D/V-AATAT_9_* - 5.26%. We speculated that the mild phenotype could either be due to the absence of D1 protein in the DV border cells or its low expression. Therefore, we overexpressed D1 protein using *C96Gal4,* which expresses the protein in the DV border cells, where the D/V-enhancer is active (Sup. Figure 3F). Overexpression of D1 in the homozygous *D/V-SSR* lines resulted in a range of phenotypes: T1 - wild type, T2 - has less than three notches or loss of bristles or both, T3 - has three or more notches, T4 - decreased wing length with disorganized arrangement of cells and T5 - small sized wings (Figure 4B). Upon D1 overexpression, the percentage of T1 wings was highest for *D/V-AAAAT_8_* at 10% and the least for *D/V-AAAT_10_* at 0.38%, with the order being *D/V- AAAAT_8_*>*D/V-AAT_14_*>*D/V-AATAT_9_*>*D/V-AAAT_10_*. The T4 and T5 phenotypes upon D1 overexpression were highest for *D/V-AAAT_10_* and the lowest for *D/V-AAT_14_*, with the order being *D/V-AAAT_10_*∼*D/V-AATAT_9_*>*D/V-AAAAT_8_*>*D/V-AAT_14_* (Figure 4C). At the same time, the T2 phenotype, which was the milder phenotype, was the highest for *D/V-AAT_14_*, with the order being *D/V-AAT_14_*>*D/V-AAAAT_8_*>*D/V-AATAT_9_*>*D/V-AAAT_10_*, which is the reverse order of extreme phenotypes (Figure 4C). However, overexpression of D1 without the SSR background (*C96Gal4>UASD1*) majorly resulted in only two phenotypes, T2 and T3, with the percentage of T3 being at least 75%. We did observe T4 and T5, but they were less than 1% wherever observed (Figure 4C). These results indicate that AAAT_10_ and AATAT_9_ function as strong enhancer blockers, whereas AAT_14_ functions as the weakest among the four AT-rich SSRs.

**Figure 4:**
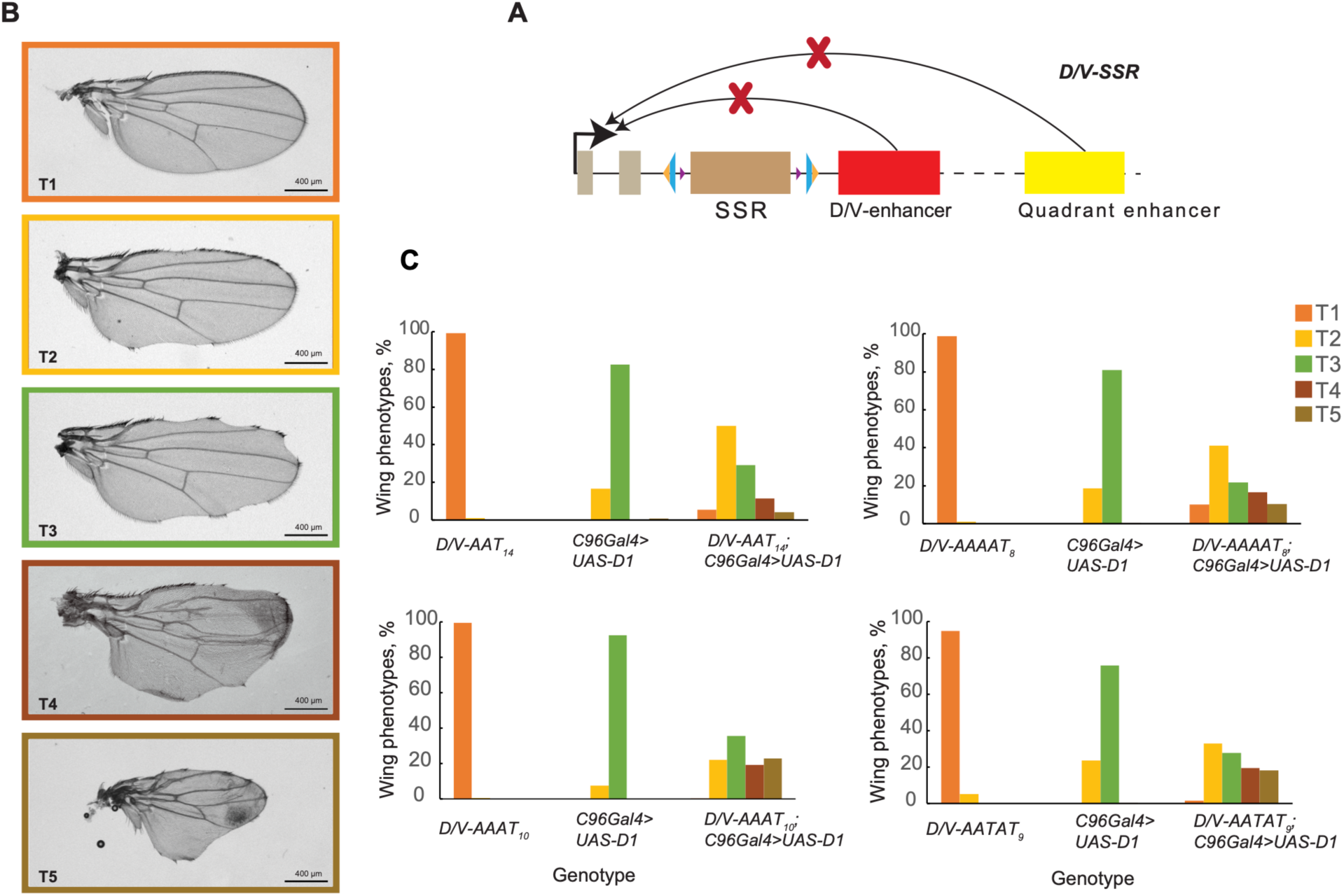
AT-rich SSRs function as enhancer blockers upon overexpression of D1 in the DV border cells using C96Gal4. **A)** Schematic showing the SSR integrated *ON-enhancer attP* line. *loxP* sites (violet triangles) are depicted. **B)** Wing phenotypes observed upon overexpression of D1 in the homozygous *DN-SSR* background: T1 - wildtype wings; T2-loss of bristles or <3 notches or both; T3-more than 2 notches; T4-decrease in wing length with disorganized wing cells; TS-decreased size of the wings. **C)** Percentage of different wing phenotypes across various genotypes: *DN-SSR, C96Ga/4>UASD1* was the control and D1 overexpression in the homozygous *DN-SSR* lines *(DN-SSR; C96Ga/4>UASD1)*.

## Discussion

SSRs regulate multiple biological processes (Horton et al. 2023; Gymrek et al. 2016; Hui et al. 2003; Hammock and Young 2005) and uncontrolled expansion of certain SSRs results in repeat expansion disorders (Depienne and Mandel 2021). However, their normal physiological roles are poorly understood. Genomic analysis of 719 organisms across the animal kingdom showed that 26% of 501 SSRs show length-specific enrichment in at least one organism (Srivastava et al. 2019). Several human SSRs enriched at longer repeat lengths were shown to function as enhancers or silencers, enhancer blockers and barrier elements in the K562 cell line (Krishnan et al. 2017). We sought to investigate the mechanistic basis of the cis-regulatory functions of these SSRs. We show that SSRs bind proteins in a sequence-specific manner and identified 33 unique SSR-binding proteins known to be involved in transcription, heterochromatin binding, DNA repair and RNA splicing enriched in 8 SSR pull-downs. One of these proteins, D1, was enriched in 6 SSRs-AAT_14_, AAAT_10_, AAAAT_8_, AATAT_9_, AAAG_13_ and AAAAG_11_. Further, using a novel *in vivo* enhancer blocker assay in *Drosophila*, we demonstrate that 4 AT-rich SSRs function as enhancer blockers in the presence of D1 protein.

The first enhancer blocker, *gypsy* retrotransposon was discovered by characterizing the *yellow* gene allele, *y[2]* (Geyer and Corces 1992). The *yellow* gene is regulated by multiple enhancers, which bring about wild-type pigmentation in various body parts. In the *y[2]* allele, *gypsy* blocks only wing blade and adult cuticle enhancers from interacting with *yellow* promoter while other *yellow* gene enhancers are unaffected. This results in yellow pigmentation of wing blades and adult cuticle only (Geyer and Corces 1992). The *y[2]* allele was extensively used to characterize *gypsy*’s regulators such as Mod(mdg4) (Gerasimova et al. 1995), CP190 (Pai et al. 2004), Polycomb and Trithorax group of proteins (Gerasimova and Corces 1998), dTopors (Capelson and Corces 2005), SUMO conjugation pathway members (Capelson and Corces 2006) and the RNAi machinery (Lei and Corces 2006).

Multiple P-element-based enhancer blocker assays were developed by Paul Schedl and others in *Drosophila* to test new enhancer blockers (Kellum and Schedl 1992; Vazquez and Schedl 1994; Hagstrom et al. 1996; Cai and Levine 1995). These include the *yp1* enhancer-based assay regulating *hsp70:lacZ* in fatbody (Kellum and Schedl 1992); the *ftz* enhancers-based assay that regulates *hsp70:lacZ* in embryonic stages and in the central nervous system of germband extended embryos (Hagstrom et al. 1996); and the *white* enhancer-based assay regulating the *mini-white* reporter gene (Vazquez and Schedl 1994). All these assays are subject to position effects and require the generation of multiple transgenic lines. Further, comparison between test elements is unreliable when using these assays. The *ftz* assay is less sensitive to position effects when compared to the *white* promoter-*mini white* assay (Barges et al. 2000). However, β-gal staining as readout in lacZ-based assays is low-throughput and incompatible with genetic screening. The *ftz* assay has been used extensively to test and confirm enhancer blockers like *fab-8* (Barges et al. 2000), ME boundary (Sultana et al. 2011), EB boundary (Verma et al. 2021), GATA SSR (Kumar et al. 2013), GA SSR (Vasanthi et al. 2010) and enhancer blockers from *Anopheles gambiae* (Ahanger et al. 2013). Compared to the *y[2]* allele, fewer studies have attempted to use *ftz* and *miniwhite* assays to identify the regulators of enhancer blockers.

Overall, the traditional enhancer blocker assays suffer from position effects, have low throughput due to cumbersome molecular assays required to assess phenotype and are not feasible for traditional genetic screens (Vazquez and Schedl 1994; Kellum and Schedl 1992; Hagstrom et al. 1996). To overcome these drawbacks, we designed a novel enhancer blocker assay in *Drosophila* using the *vestigial* locus. The *vestigial* gene is responsible for wing and haltere development and is regulated by D/V and quadrant enhancers (Kim et al. 1996). In our assay, blocking the D/V enhancer activity gives rise to wing and haltere phenotypes, which are visually detectable. It utilizes the attP-attB based recombination for site-specific integration of test fragments, which helps avoid position effects and makes comparisons of different test enhancer blockers feasible. It allows for studying the enhancer blocking activity at an endogenous locus rather than in the transgenic context. It also allows estimation of enhancer blocker activity by direct quantification of wing phenotypes without the need for molecular biological assays. Moreover, it is amenable to genetic screens to identify the various regulators of these tested enhancer blockers. Our assay can be used to characterize enhancer blockers and their regulators *in vivo*.

Analysis with *gypsy*, a known enhancer blocker, confirmed the validity of our assay. *D/V-gypsy* flies show a range of phenotypes characteristic of *vestigial* loss of function, such as loss of one or both wings and/or halteres, upright post-scutellar bristles and lethality (Figure 3C and Sup. Figure 3C-3D). These phenotypes were rescued upon *su(Hw)* or *mod(mdg4)* or *Cp190* mutations as *gypsy* enhancer blocker activity is mediated by a complex of proteins consisting of Su(Hw), Mod(Mdg4) and CP190 (Figure 3E-G).

We proceeded to confirm the cis-regulatory functions of SSRs *in vivo* using our assay. Since D1 protein was enriched in multiple different AT-rich SSRs, we investigated if D1, along with AT-rich SSRs, functions as enhancer blockers *in vivo*. We generated 4 transgenic lines with different AT-rich SSRs. Integration of SSRs resulted in a mild phenotype in less than 5% of the flies. However, overexpression of D1 protein using *C96Gal4* in the DV border cells of the wing pouch of the homozygous *D/V-SSR* lines resulted in wing phenotypes of varying degrees. The order of enhancer blocking activity observed was AAAT_10_∼AATAT_9_>AAAAT_8_>AAT_14_ (Figure 4). Thus, we show for the first time that AT-rich SSRs, along with D1 protein, function as enhancer blockers *in vivo*. We identified several candidate proteins and SSRs that may be analyzed using our enhancer blocker assay in the future.

In our pull-downs followed by proteomics experiments, we identified 33 unique proteins that bind SSRs. Some of these proteins bind more than one SSR. Proteins like D1, Jigr1 and Rrp1 bound several SSRs in our proteomic analysis (Sup. Table 2). It has been shown that D1 or Jigr1 when overexpressed in combination with Jing in the eyes of *Drosophila*, results in eye defects (Sun et al. 2006). The roles of D1 or Jigr1 individually or together as part of a complex, along with AT-SSRs, may be interesting to investigate further. Similarly, 6 of the 9 proteins that enriched in AATAT_9_ also enriched in AAAAT_8_. We speculate that proteins that bind multiple SSRs bind them with differing affinities and can coregulate genes with these SSRs in their regulatory regions.

Several TFs that bound SSRs have the MADF (myb/SANT-like domain in Adf-1) domain. For example, Jigr1 enriched in AAT_14_, AAAT_10_, AAAAT_8_ and AATAT_9_ SSRs; CG8944 enriched in AATAT_9_ SSR; Dip3 enriched in AAAAG_11_ and AAAG_13_ and Prod, CG14005 and CG7239 enriched in ACATAT_8_ SSRs. The myb homology domain, which is similar to the MADF domain, is present in TRF1 and TRF2, the mammalian telomere SSR-(TTAGGG) binding proteins, and is required for DNA binding (Bianchi et al. 1997; England et al. 1992). We think that the MADF domain could be a common SSR-binding protein domain. Further, MADF protein Dip3 has a BESS domain involved in protein-protein interactions. MADF-BESS domain-containing proteins are known to associate with heterochromatin and regulate transcription (Cutler et al. 1998; Shukla et al. 2014). Recently, Mzfp1 was shown to bind promoters and is required for insulator activity and housekeeping gene expression. Mzfp1 contains 2 MADF domains and 6C2H2 Zn fingers (Sokolov et al. 2024). CG8944 enriched in AATAT binding proteins has multiple C2H2 Zinc fingers along with 2 MADF domains. It may be possible that SSRs, along with their MADF domain-containing binding proteins, may function as enhancer blockers. This can be investigated using our assay in the future.

Taken together, in this study, we identified several SSR binding proteins and developed a novel *vestigial* enhancer blocker assay that can be used to test cis-regulatory functions of SSRs and their binding proteins. Our assay utilizes an endogenous locus that is easily scorable, free of position effects and accessible to high throughput traditional genetic approach. Using this assay, we demonstrate that AT-rich SSRs AAT_14_, AAAT_10_, AAAAT_8_ and AATAT_9_ are capable of enhancer blocking activity in conjunction with their binding protein, D1, *in vivo*, in *Drosophila*. Our assay provides an easier and unique tool to not only identify the enhancer blocker function of SSRs and their binding proteins but also other enhancer blockers and their regulators.

## Materials and Methods

### Preparation of nuclear extract from *Drosophila* embryos

10 g of dechorionated 0-16 h *Drosophila* embryos were suspended in 20 ml nuclei isolation buffer I (NIB I) (15 mM HEPES, pH 7.6, 40 mM KCl, 1 mM EDTA, 0.25 M sucrose, 0.1 mM spermine, 0.125 mM spermidine, PMSF, 1X protease inhibitor cocktail (PIC)) and homogenized until embryos were disrupted. The homogenized embryos were filtered through Mira cloth and washed with 5 ml of NIB I. The filtrate was centrifuged at 600 g for 30 s at 4°C. The supernatant was mixed with an equal volume of NIB II (15 mM HEPES, pH 7.6, 40 mM KCl, 1 mM EDTA, 1.8 M sucrose, 0.1 mM spermine, 0.125 mM spermidine, PMSF, 1X PIC) and centrifuged at 8000 rpm at 4°C. The nuclear pellet obtained was submerged in NIB I for 5 min on ice, resuspended and centrifuged at 1000 g for 10 minutes at 4°C. The above step was repeated again, and the nuclear pellet was resuspended in 2 ml of NEB 20 (10 mM HEPES, pH 7.6, 20 mM KCl, 3 mM MgCl_2_, 1 mM EDTA, 1M sucrose, 10% glycerol, 1X PIC, PMSF). To this, an equal volume of NEB 700 (10 mM HEPES, pH 7.6, 700 mM KCl, 3 mM MgCl_2_, 1 mM EDTA, 1 M sucrose, 10% glycerol, 1X PIC, PMSF) was added slowly with mixing. The solution was rocked gently for 1 h at 4°C. Subsequently, it was centrifuged at 42,000 rpm for 1 h. The supernatant (nuclear extract) was collected and stored as aliquots at -80°C.

### Electrophoretic Mobility Shift Assay (EMSA)

Synthesized oligos were annealed in an annealing buffer (10mM Tris, pH 8, 50mM NaCl, 1mM EDTA) using the following program on a thermocycler: 95° C for 2 min, cooling -2° C/s to 84° C, and -0.1° C/s to 25° C. 15 pmol of annealed oligos were end-labeled using **γ**^32^P-ATP and T4 Polynucleotide kinase (New England Biolabs) for 1 h at 37° C. The radiolabeled oligos were purified using Sephadex G-50 (Sigma) column. The eluate was adjusted to 150 μl so that the final concentration of the oligos was 0.1 pmol/μl.

Binding reactions were performed in a reaction volume of 15 μl containing 25 mM HEPES, pH 7.6, 10% glycerol, 1.5 μg tRNA, 1 mM EDTA, 150 mM NaCl (total), 0.3 pmol of radiolabeled oligos and 4 μg nuclear extract. The binding reactions were incubated at RT for 15 mins, followed by incubation at 4° C for another 15 mins. For competition reactions, unlabeled oligos were added to the binding reactions at either 10-fold, 20-fold, or 50-fold higher concentrations compared to the radiolabeled oligos. Following incubation, the binding reaction mixtures were loaded onto a 4% acrylamide-bisacrylamide gel (39:1 acrylamide: bisacrylamide) in 0.5X Tris-borate-EDTA (TBE), and run at 15 mA at 4°C for 3 h in 0.5X TBE buffer. The gels were subsequently dried and exposed on a phosphor screen, which was imaged using a phosphorimager.

### DNA-based protein pulldowns

SSR DNA was amplified from SSRs in *pGL3* vectors (Krishnan et al. 2017) using a 5’ biotinylated forward primer and an unmodified reverse primer. The control DNA was amplified using the same set of primers from the *pGL3* vector without the SSRs. 5 μg of purified biotinylated DNA was bound to 20 μl M280 streptavidin beads in 1X binding buffer (5 mM Tris, pH 7.5, 0.5 mM EDTA, 1 M NaCl) at 4° C. The beads were washed twice with 1X binding buffer and collected using a magnetic stand. 20 μl of DNA bound beads were incubated with 200 μg of 0-16 h *Drosophila* embryo nuclear extract in 25 mM HEPES, pH 7.6, 10% glycerol, 1.5 μg tRNA, 1 mM EDTA, 150 mM NaCl, 1X PIC. The protein-bound beads were separated and washed 3x for 10 min at 4° C with wash buffer (25 mM HEPES, pH 7.6, 10% glycerol, 1.5 μg tRNA, 1 mM EDTA, 150 mM NaCl, 1X PIC). The beads were then boiled in 1X Laemlli buffer for 5 min at 95° C, the supernatant was collected and stored at -80° C by separating the beads using a magnetic stand. The protein pull-downs were performed in sets of 2 SSRs and a control, in biological triplicates. In total, we performed eight SSR and four control pull-downs in triplicates, amounting to 36 MS-MS runs.

### In-Gel digestion for LC-MS/MS

The pulled-down protein mixture was resolved for about 1 cm on 12% SDS-PAGE. The gel was cut into 1 mm³ pieces and washed thrice in 50% ACN in 25 mM NH_4_HCO_3_, followed by dehydration in 100% ACN with vigorous vortexing. Dehydrated gel pieces were vacuum dried for 20 min, followed by reduction of disulfide bonds in 10 mM DTT in 25 mM NH_4_HCO_3_ at 55°C for 45 min. Subsequently, the gel pieces were incubated in 55 mM Iodoacetamide in 25 mM NH_4_HCO_3_ at RT in the dark for 45 min. Then the gel pieces were dehydrated in 100% ACN and vacuum dried for 10 min. The dried gel pieces were incubated with trypsin gold at 37°C for 16 h. The tryptic peptides were eluted twice in 2.5% TFA in 50% ACN by incubating the gel pieces at RT with vigorous vortexing for 30 mins. The supernatant containing peptides was quickly vacuum dried and resuspended in 2% ACN in 0.1% formic acid. The peptides were desalted using p10 Ziptip C18 columns and the desalted peptides were resuspended in 2% ACN in 0.1% formic acid.

### Proteomics and data analysis

Proteomic runs were performed on Thermoscientific Q-Exactive HF using a 60 min acetonitrile gradient. The spectra obtained were analyzed using MaxQuant version 2.1.0.0 with default parameters, except that “Label-free quantification” and “match between runs” parameters were applied (Cox and Mann 2008). The protein groups files obtained from MaxQuant were further analyzed using LFQ analyst at default parameters (Shah et al. 2020). Volcano plots were generated using VolcaNoseR with the output table obtained from LFQ analyst (Goedhart and Luijsterburg 2020).

### Molecular cloning

To generate gRNA plasmids, annealed gRNA oligos were ligated into *BbsI* digested *pCFD3*. To generate donor plasmids, *SmaI* digested *pBSKS* was ligated with *attP1_220__3XP3GFP*. The resultant plasmid was digested with *SacI* and *SpeI* and ligated with left homology arms. The left homology arms for the *D/V-enhancer attP* line were amplified by LfDV_F and LfDV_R, and the *QE-enhancer attP* line was amplified by LfQE_F and LFQE_R. The generated plasmid was subsequently digested with *SalI* and *KpnI* and ligated with the right homology arms. The right homology arms were amplified with RtDV_F and RtDV_R for the *D/V-enhancer attP* line and RtQE_F and RtQE_R for the *QE-enhancer attP* line. The resultant plasmids were digested with *EcoRV* and ligated with *attP2_220_* (orientation opposite to attP1) for both vectors. For the *pBSKS_attB1_40__attB2_40_* plasmid, attB_40_ oligos were annealed and ligated into *SacI* digested *pBSKS* and the resulting plasmid was digested with *KpnI* and ligated with another set of attB_40_ oligos in opposite orientation to generate *pBSKS_attB1_40__attB2_40_* plasmid. *gypsy* was cut from *pCFD3* plasmid using *StuI* and *BglII sites* and ligated into *SmaI* and *BglII* digested *LML* (*LoxP1-MCS-LoxP2*) vector (Vasanthi et al. 2010). *SmaI* digested *LML* vector was ligated with annealed oligos *AAT_14_* or *AAAT_10_* or *AAAAT_8_* or *AATAT_9_*. Both *gypsy* or *SSRs* ligated *LML* plasmids were digested with *NotI* and the resulting *gypsy* or *SSR* fragments were ligated into *pBSKS_attB1_40__attB2_40_* plasmid using the *NotI* site.

### Injection and screening of *attP* lines

All embryo injections were performed at C-CAMP (Centre for Cellular and Molecular Platforms), Bengaluru, India. *nanos-cas9* embryos (Bloomington St. No. 54591) were injected with 250 ng/μl of *pCFD3* vector and 500 ng/μl of donor vector. Injected *nanos-Cas9* flies were mated to *w; Pin/CyO* flies in single fly crosses. The GFP-positive and *CyO* flies were subsequently crossed to *w; Pin/CyO* flies again in single fly crosses. The GFP-positive and *CyO*-only flies in the next generation were selfed, the lines balanced and sequence confirmed. Eventually, the *nanos-Cas9* containing X chromosome was crossed out if present.

### Injection and screening for *gypsy* or *SSR* integrated flies

500 ng/μl of the *pBSKS_attB1_40_-test-attB2_40_* plasmids were injected into *nanos-PhiC31 integrase; attP1_220_-3XP3GFP-attP2_220_* embryos. Injected *nanos-PhiC31 integrase; attP1_220_-3XP3GFP-attP2_220_* male flies were crossed to *w; Pin/CyO* virgins in single fly crosses, and their male progeny with *CyO* and loss of GFP were subsequently crossed to *w; Pin/CyO* virgins in single fly crosses in the next generation. The resulting *CyO*-only flies were selfed, maintained and sequence confirmed. The *SSR* integrated flies that had forward orientation (with respect to SSR) were phenotyped and the line that showed the least phenotype was used for overexpression of D1.

### Flies used for the experiments

Genotypes of flies used in our experiments-

1. w; D/V-gypsy/CyO; Cp190[H4-1]/TM6B 2. w; D/V-gypsy/CyO; mod(mdg4)[ul]/TM6B 3. y[2] sc[1] v[1]/ w ; D/V-gypsy/CyO; Su(Hw)[8]/TM6B 4. w; D/V-SSR/CyO; UASD1/TM2 or TM6B 5. w; D/V-SSR/CyO; C96-Gal4/TM2 or TM6B

Bloomington stock (Bl.S.) No. 59960 (*mod(mdg4)*), Bl. S. No. 59965 (*Cp190*), Bl. S. No. 1053 (*su(Hw)*), Bl. S. No. 25757 (*C96Gal4*), fly orf Stock No. F000455 (*UAS-D1-3XHA*) fly stocks were used to make the above flies.

### Phenotyping wings

*w; D/V-SSR; UASD1/C96Gal4* were obtained by crossing *w; D/V-SSR; UASD1/UASD1* with *w; D/V-SSR; C96Gal4/TM2*. A minimum of 200 wings (95 wings for *D/V-AATAT_9_*) of the controls and test flies were phenotyped according to the phenotyping scale shown in Figure 4B. The wing images were taken using Axio Zoom.V16 after dissecting and mounting the wings from female flies.

## Acknowledgments

We thank V. Bharathi for wing imaginal disc dissections and immunostaining, V. Bharathi, Sonal Nagarkar Jaiswal and the past and present members of the RKM lab for their constructive suggestions.

## Data Availability

Mass spectrometry proteomics data are deposited to the ProteomeXchange Consortium via the PRIDE partner repository with dataset identifier PXD058086.

## Author Contributions

Designed experiments: K.P., R. K. M.; Performed experiments: K.P., F.A.; Data analysis: K. P., R.K.M.; Wrote manuscript: K.P., F.A., R.K.M.

