## Supplementary material for "A novel enhancer blocker assay identifies AT-rich simple sequence repeats as D1-dependent enhancer blockers in *Drosophila*": Sup. Tables 1-3

**Supplementary Table 1**: List of enriched proteins across various SSRs whose $log_2$(foldchange over control) > 1.5 and $-log_{10}$(adj. p-value) > 2

| SSR | SSR binding proteins | | | | | |
|---|---|---|---|---|---|---|
| **$AAAAG_{11}$** | D1 | Dip3 | Rrp1.1 | | | |
| **$AAAAT_8$** | D1 | Jigr1 | Phr (Photorepair) | Phr6-4 (Photolyase) | Rox8 | Jarid2 | Blm |
| | Tet | | | | | | |
| **$AAAT_{10}$** | D1 | Jigr1 | Q9VXL5 (CG8924/CC9) | | | | |
| **$AAT_{14}$** | D1 | Jigr1 | Q9VXL5 (CG8924/CC9) | Mef2-RC | | | |
| **$AATAT_9$** | D1 | Jigr1 | phr | Phr6-4 | Tet | Jarid2 | Abd-B |
| | Top1 | Q9VXQ6 (CG8944) | | | | | |
| **$ACATAT_8$** | Cf2 | Q9VMM3 (CG7239) | Top2 | Q7JWH6 (CG1888) | prod | Ibf1 | M9PB45 (CG14005) |
| | Rrp1.1 | cg | | | | | |
| **$AGAT_{10}$** | arm | Diminutive (dm) | Cutlet | x16 | Rrp1.1 | Moca-cyp | A0A0B4LFE1 (CG10139) |
| **$AAAG_{13}$** | Dip3 | D1 | lid | Rrp1.1 | x16 | Top1 | Diminutive (dm) |
| | CG1815 (Zmynd 8) | Q7K4L8 (CG7878) | Jupiter | | | | |

**Supplementary Table 2**: Enriched proteins identified in more than one SSR pull down

| **Gene names** | **No. of SSRs enriched in** | **SSRs in which enriched** |
|---|---|---|
| D1 | 6 | $AAT_{14}$, $AAAT_{10}$, $AAAAT_{8}$, $AATAT_{9}$, $AAAAG_{11}$, $AAAG_{13}$ |
| Jigr1 | 4 | $AAT_{14}$, $AAAT_{10}$, $AAAAT_{8}$, $AATAT_{9}$ |
| phr | 2 | $AAAAT_{8}$, $AATAT_{9}$ |
| Dip3 | 2 | $AAAG_{13}$, $AAAAG_{11}$ |
| Jarid2 | 2 | $AAAAT_{8}$, $AATAT_{9}$ |
| Phr6-4 | 2 | $AAAAT_{8}$, $AATAT_{9}$ |
| Rrp1 | 4 | $AAAAG_{11}$, $ACATAT_{8}$, $AAAG_{13}$, $AGAT_{10}$ |
| CC9/Q9VXL5/CG8924 | 2 | $AAT_{14}$, $AAAT_{10}$ |
| Tet | 2 | $AAAAT_{8}$, $AATAT_{9}$ |
| x16 | 2 | $AAAG_{13}$, $AGAT_{10}$ |
| dm | 2 | $AAAG_{13}$, $AGAT_{10}$ |
| Top1 | 2 | $AAAG_{13}$, $AATAT_{9}$ |

**Supplementary Table 3:** Primers used in this study

| Primer Name | Sequence |
|---|---|
| D/V-Enhancer_gRNA_sense | GTCGTCTCAGCGAAATCTCTGTAG |
| D/V-Enhancer_gRNA_antisense | AAACCTACAGAGATTTCGCTGAGA |
| QEnhancer_gRNA_sense | GTCGATTCATAAGTGGATATTCC |
| QEnhancer_gRNA_antisense | AAACGGAATATCCACTTATGAATC |
| LfDV_F | GTAGAGCTCTCCAAGTTTTAGCCCCCTTC |
| LfDV_R | GGACTAGTCAGAGATTTCGCTGAGATTTTGA |
| LfQE_F | GGAGAGCTCAAGCGAAAAGTTTTGGGGCT |
| LfQE_R | CCGACTAGTTCCCGGCAGATTCCCGAAAG |
| RtDV_F | GGAGTCGACTAGTGGCAATCACAATTTCTCTT |
| RtDV_R | GGGGTACCAACCAGCGATCGTGAGAACT |
| RtQE_F | GGAGTCGACATATCCACTTATGAATCCAG |
| RtQE_R | GGGGTACCTCAGACAGGGCGAGAGAAAA |
| attP_F | AGTACTGACGGACACACCGAA |
| attP_R | TCGCGCTCGCGCGACTGACG |
| 3XP3GFP_F | CCGGGGATCTAATTCAATTAGAG |
| 3XP3GFP_R | CTAAGATACATTGATGAGTTTGG |
| attB40_SacI_F | CCGGGTGCCAGGGCGTGCCCTTGGGCTCCCCG GGCGCGTACGAGCT |
| attB40_SacI_R | CGTACGCGCCCGGGGAGCCCAAGGGCACGCCC TGGCACCCGGAGCT |
| attB40_KpnI_F | CGTACGCGCCCGGGGAGCCCAAGGGCACGCCC TGGCACCCGGGTAC |
| attB40_KpnI_R | CCGGGTGCCAGGGCGTGCCCTTGGGCTCCCCG GGCGCGTACGGTAC |
| Npfp | CCGAGCTCTTACGCGTGC |
| Pgl3ssrR1a | GGTTGCTGACTAATTGAGATGC |
