## Supplementary material for "A novel enhancer blocker assay identifies AT-rich simple sequence repeats as D1-dependent enhancer blockers in *Drosophila*": Sup. Figure 1

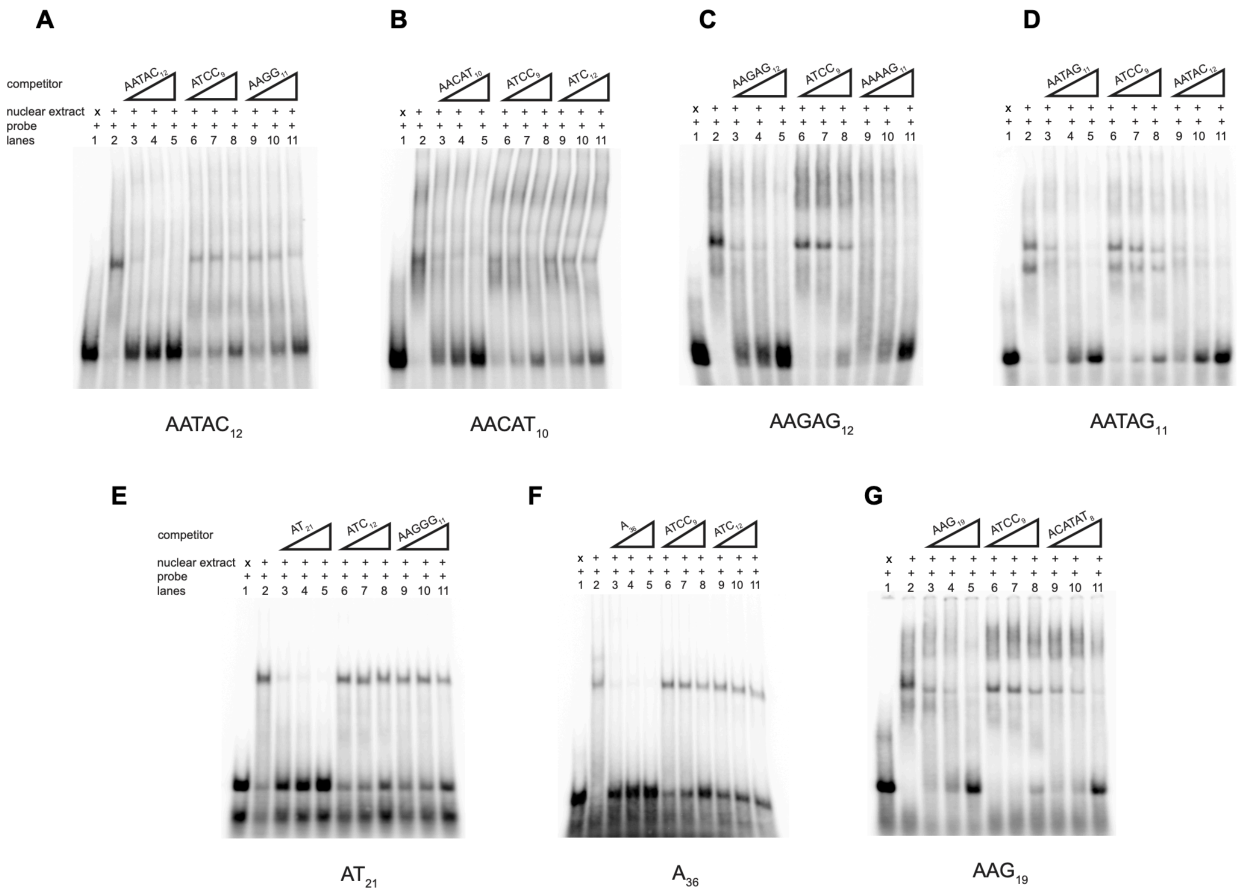


**Sup. Figure 1: Electrophoretic mobility shift assays of length-enriched SSRs in the presence of nuclear extract from 0-16 h *Drosophila* embryos.** Each gel shows DNA-protein complexes of radiolabeled SSRs in lane 2. Competition reactions with the same but unlabeled SSR are shown at 10- or 20- or 50-fold concentrations higher than radiolabeled probe in lanes 3-5 of each gel. Competition reactions with a different unlabeled SSR dissimilar in sequence to the radiolabeled probes are shown at 10- or 20- or 50-fold concentrations in lanes 6-8 of all gels, lanes 9-11 of A, B and E-G. Competition reactions with a different unlabeled SSR with sequence similarity to the radiolabeled probes are shown at 10- or 20- or 50-fold concentrations in lanes 9-11 of gels C-D.
