## Supplementary material for "A novel enhancer blocker assay identifies AT-rich simple sequence repeats as D1-dependent enhancer blockers in *Drosophila*": Sup. Figure 2

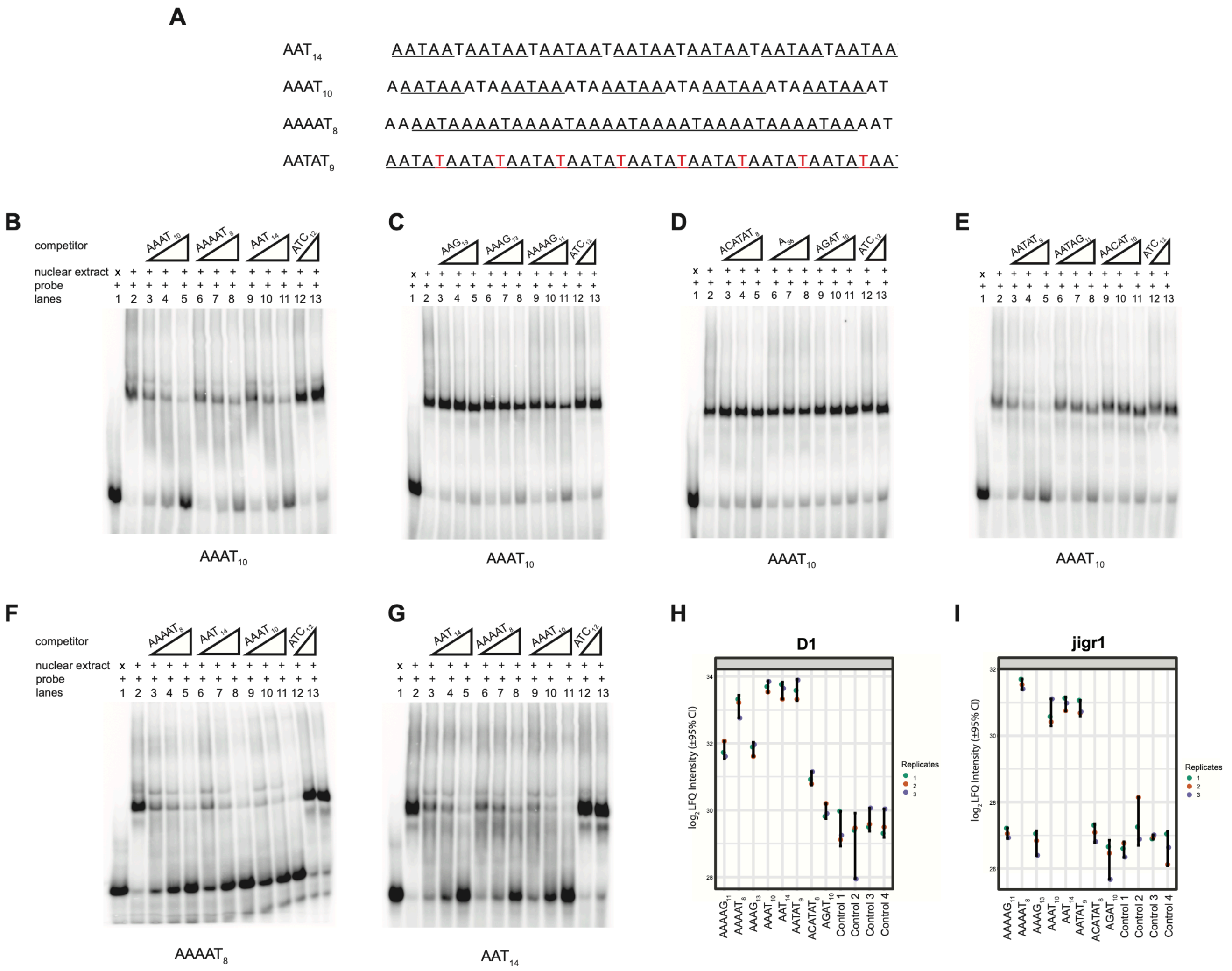


**Sup. Figure 2: A)** Alignment of $AAT_{14}$, $AAAT_{10}$, $AAAAT_8$ and $AATAT_9$ shows the recurrence of AATAA motifs, see underlined. The motifs of $AATAT_9$, AATAT, differ from that of AATAA in the last nucleotide which is highlighted in red. **B-E)** Competition EMSAs of $AAAT_{10}$ radiolabeled probe: EMSAs with radiolabeled $AAAT_{10}$ using nuclear extract from 0-16 h *Drosophila* embryos. Lanes 3-5, 6-8, 9-11 depict competition of shifts with various unlabeled SSRs at higher concentrations of 10 or 20 or 50-fold compared to the radiolabeled $AAAT_{10}$. All gels show competition with unlabeled $ATC_{12}$ performed at 10 and 50-fold compared to radiolabeled $AAAT_{10}$. **F-G)** Competition EMSAs of $AAAAT_8$ and $AAT_{14}$ radiolabeled probe: EMSAs with radiolabeled probe using nuclear extract from 0-16 h *Drosophila* embryos. Lanes 3-11 depict competition with unlabeled $AAT_{14}$, $AAAT_{10}$ and $AAAAT_8$ at higher concentrations of 10 or 20 or 50-fold compared to the radiolabeled probe. Competition with unlabeled $ATC_{12}$ was performed at 10 and 50-fold compared to radiolabeled probe in lanes 12-13.
**H, I)** Plots showing LFQ intensities of D1 and Jigr1 across various SSR-based protein pull-downs.
