## Supplementary material for "A novel enhancer blocker assay identifies AT-rich simple sequence repeats as D1-dependent enhancer blockers in *Drosophila*": Sup. Figure 3

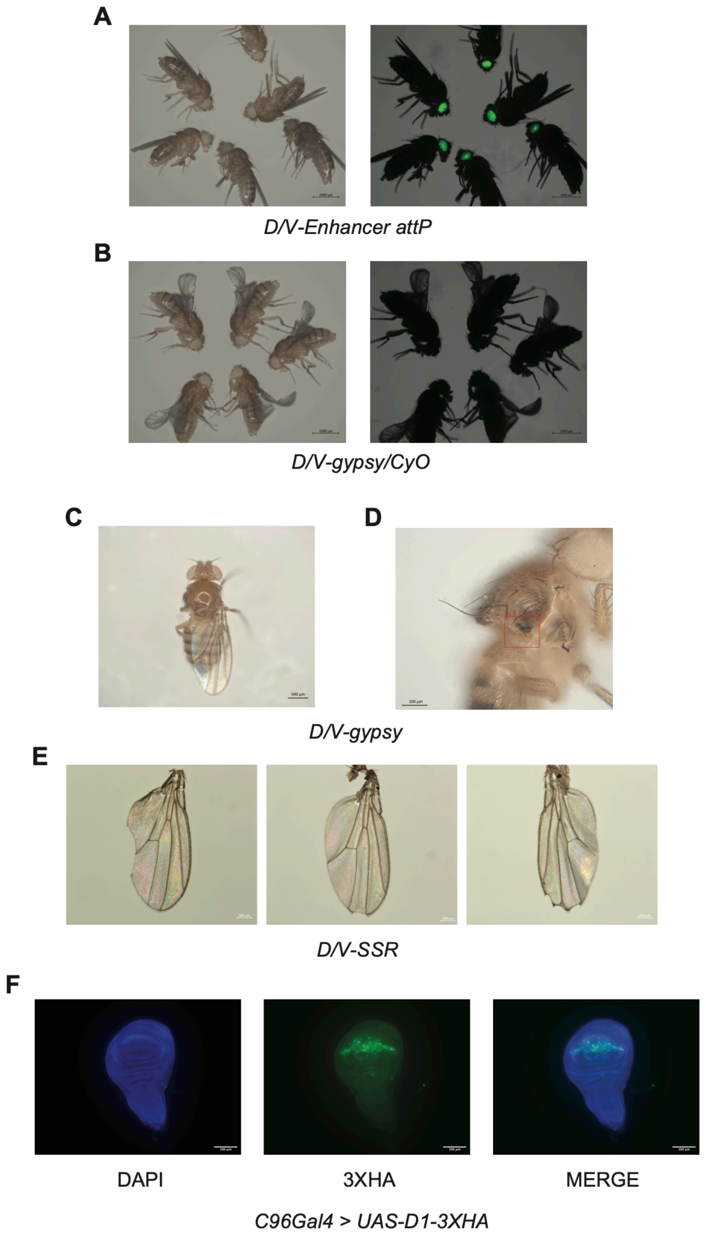


**Sup. Figure 3: A)** *D/V-Enhancer attP* flies showing the presence of GFP in the eyes. **B)** *D/V-gyspy/CyO* flies showing the loss of GFP in the eyes after RMCE. **C)** *D/V-gypsy* fly showing loss of one wing. **D)** *D/V-gypsy* fly showing upright post-scutellar bristles and the transformation of wing to notum (red box). **E)** Mild wing phenotypes observed in *D/V-SSR* flies. **F)** D1 overexpression in the DV-border cells of the wing imaginal discs of *C96Gal4>UASD1* flies.
